# Adaptive Expansion of Taste Neuron Response Profiles by Congruent Aroma in *Drosophila*

**DOI:** 10.1101/859223

**Authors:** Yiwen Zhang, Yuhan Pu, Yan Zhang, Haini N. Cai, Ping Shen

**Affiliations:** Department of Cellular Biology and Biomedical and Health Science Institute, University of Georgia, 500 D.W. Brooks Drive, Athens, GA, 30602, USA

## Abstract

Pairing of food aroma with selected taste can lead to enhanced food flavor and eating euphoria, but how cross-modal sensory combinations are integrated to increase food reward value remains largely unclear. Here we report that combined stimulation by food aroma and taste drastically increased appetite in well-nourished *Drosophila* larvae, and the appetizing effect involves a previously uncharacterized smell-taste integration process at axon terminals of two Gr43a gustatory neurons. Molecular genetic analyses of the smell-taste integration reveal a G protein-mediated tuning mechanism in two central neuropeptide F (NPF) neurons. This mechanism converts selected odor stimuli to NPF-encoded appetizing signals that potentiate Gr43a neuronal response to otherwise non-stimulating glucose or oleic acid. Further, NPF-potentiated responses to glucose and oleic acid require a Gr43a-independent and Gr43a-dependent pathway, respectively. Our finding of adaptive expansion of taste neuron response profiles by congruent aroma reveals a previously uncharacterized layer of neural complexity in food flavor perception.

## Introduction

Food flavor is primarily determined by smell and taste sensations. Pairing of congruent aroma with selected taste can lead to drastic enhancement of appetite and overeating in humans and animals alike^1–8^. These observations have raised two major questions: what are the molecular and neural underpinnings of the congruency of aroma-taste pairs, and how integrated representations of food flavors acquire various reward values. However, at present, our knowledge about these questions remains rather limited in any organism.

Appetizing aroma such as a banana-like scent can rapidly induce an aroused appetitive state in well-nourished *Drosophila* larvae, as evidenced by a persistent increase in their feeding activity in readily accessible palatable media^9^. The genetically tractable fly larva provides several advantages for the anatomical and functional analyses of neural processing of food smell and taste. The fly larva has a highly evolved yet numerically simpler central nervous system (CNS), and its brain shares organizational features similar to those of a mammalian brain^10–13^. Furthermore, the olfactory and gustatory systems of fly larvae are numerically simpler than those of adult flies^14–17^, providing a useful platform for the neural circuitry analysis of potential interactions between congruent aroma and taste.

In *Drosophila* larvae, perception of appetizing odor stimuli requires neural processing of ascending olfactory inputs by three layers of central neurons^9,13,17^. Following initial processing by the second-order projection neurons, the olfactory outputs are further processed in an assembly of four third-order dopamine (DA) neurons. However, the signals from the DA neurons remain not meaningful behaviorally, and a postsynaptic neuron expressing neuropeptide F (NPF, a fly homolog of mammalian neuropeptide Y) is recruited that selective assigns appetitive values to a fraction of such DA signals^9,18^. Fly larvae have also been shown to display appetitive responses to tastants including sugars, protein and salt^19–22^. Gustatory stimuli are relayed by peripheral taste neurons, located either exteriorly or interiorly, to the hindbrain-like subesophageal zone (SEZ) of fly larvae^23,24^. At present, how gustatory information is processed within the CNS and interacts with appetizing odor signals remains rarely unexplored.

In this work, we show that in well-nourished fly larvae, selective pairing of food aroma with congruent taste (e.g., glucose or oleic acid) differentially triggers impulsive-like feeding responses. We also provide molecular and cellular evidence for the cross-modal integration of glucose or oleic acid taste with congruent aroma. A pair of central NPF neurons, which defines a food reward system, harbors a G protein-mediated tuning mechanism that converts selected odor stimuli to NPF-encoded appetizing signals. These NPF signals are relayed to the larval hindbrain-like SEZ, and contribute to the food aroma-taste integration by potentiating Gr43a neuronal response to otherwise non-stimulating glucose and oleic acid. Further, the food smell-taste integration process resides at the axon terminals of the Gr43a neurons within the SEZ. Finally, the aroma-glucose and aroma-oleic acid interactions involve a shared neuronal pathway involving the NPF receptor (NPFR1) and DH44R1, a corticotropin release factor (CRF)-like receptor; However, they are differentially mediated by a Gr43a-indepedent and a Gr43a-dependent receptor pathway within the Gr43a neurons. Our finding of the adaptive expansion of taste neuron response profiles by congruent food aroma has revealed a previously uncharacterized cross-modal integration mechanism that underlies the congruency between food aroma and taste. Therefore, food taste sensation appears to be adaptively modulated, and may be much more complex than commonly recognized.

## Results

### Pairing appetizing aroma with selected sugar taste for appetite enhancement

To investigate the usefulness of the *Drosophila* larva model for studying molecular and neural basis of food flavor pairing, we tested the potential appetizing effect of a banana-like scent (e.g., pentyl acetate or PA) on odor-aroused larval feeding responses to10% glucose, 3% oleic acid or 0.5% tryptone (digested protein) media^25,26^. The feeding response can be quantified by measuring larval mouth hook contraction rate or ingestion of dyed food. For example, the PA stimulation of fed larvae triggered a two-fold 100% increase in dyed food ingestion, which corresponds to about 20% increase in the contraction rate of mouth hooks^9^. Fed larvae displayed similar baseline feeding activities in liquid media containing 10% glucose, 3% oleic acid or 0.5% tryptone. However, after briefly stimulated by 5μl of PA, larvae increased their feeding response to the glucose but not to fatty acid or tryptone food (Fig. 1A). These observations suggest that fly larvae may have evolved an innate cross-modal integration mechanism for pairing food aroma with selected taste.

**Figure 1.**
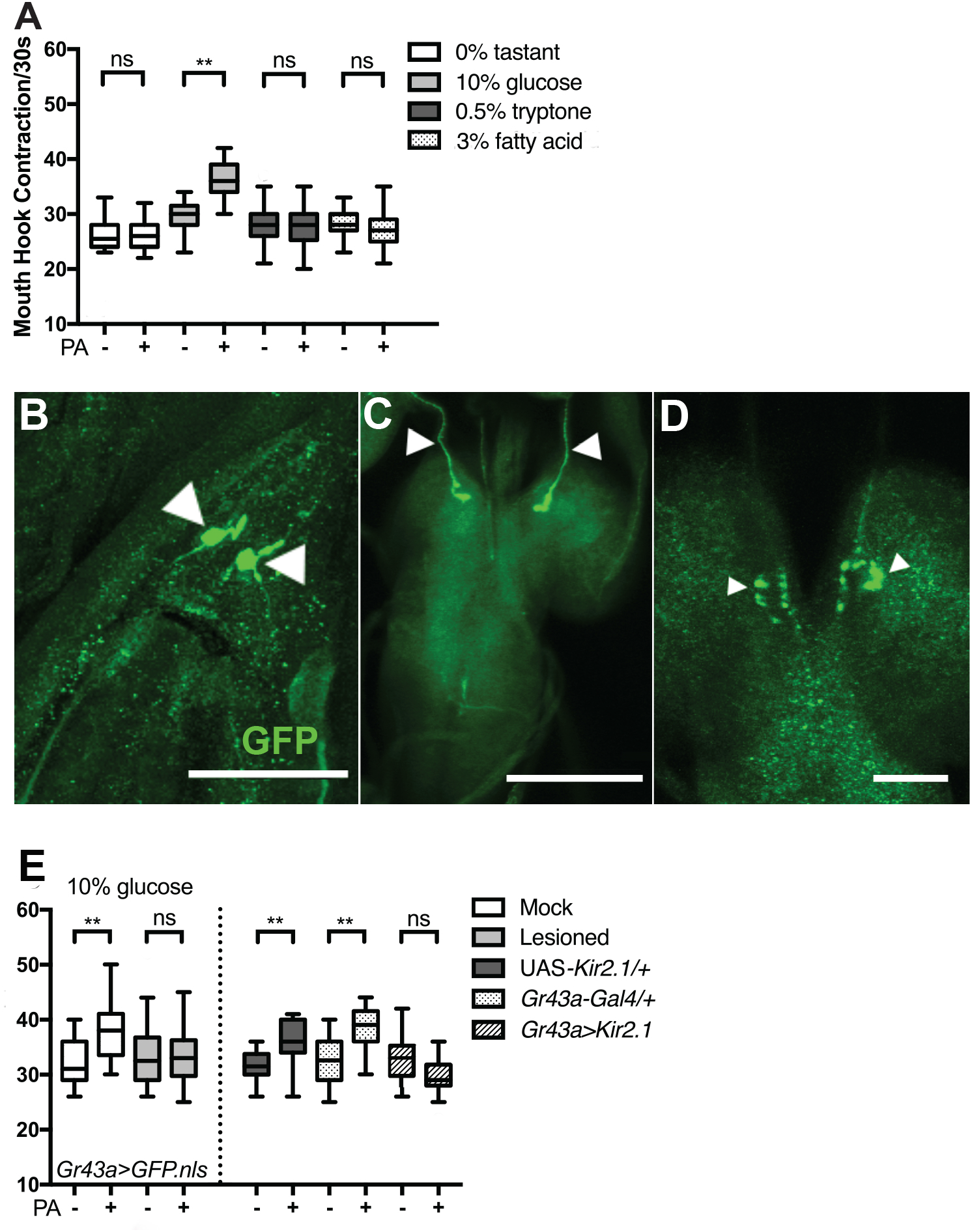
A pair of SEZ-projecting Gr43a neurons mediates the appetizing effect of food aroma and glucose taste. **(A)** Fed larvae showed similar baseline feeding activities in glucose, tryptone or fatty acid food. After stimulation by an effective dose of PA, larvae selectively increased feeding response to glucose food only. n = 17-49. Kruskal-Wallis test followed by Dunn’s multiple comparisons; ** P < 0.0001. **(B)** Lesioning of a pair of peripheral Gr43a neurons abolished PA-aroused glucose feeding. n = 14-17. One-way ANOVA followed by Sidak’s multiple comparisons test; ** P < 0.01. Kir2.1-mediated silencing of the Gr43a neurons also abolished the feeding response. n = 15-24. Kruskal-Wallis test followed by Dunn’s multiple comparisons; ** P < 0.001. **(C-E)** Expression of UAS-*mCD8∷GFP*, directed by *Gr43a-Gal4*, in early third-instar larvae: the left panel shows the soma of two peripheral Gr43a neurons at the pharyngeal region (white arrowheads). Scale bar = 60 um. The middle shows a pair of projections of the *Gr43a-Gal4* neurons innervating the SEZ (arrows). Scale bar = 100 um. The right shows a magnified view of axon terminals of the *Gr43a-Gal4* neurons at the SEZ (arrowheads). Scale bar = 25 um. All behavioral assays in this and other figures were performed under the blind condition.

### Two Gr43a neurons mediate PA-aroused glucose feeding via NPFR1

To better understand how pairing the appetizing scent with glucose taste synergistically enhances appetite, we performed a *Gal4*-based screen for candidate neurons involved in the smell-taste integration. The *Gr43a-Gal4* driver labels two gustatory neurons in the larval pharyngeal region that are responsive to fructose but not glucose stimulation^16^ (Fig. 1B-D; Fig. S1). However, to our surprise, expression of an inward-rectifying potassium channel Kir2.1 in the *Gr43a-Gal4* neurons completely abolished the PA-aroused glucose feeding. In parallel, targeted lesioning of the two *Gr43a-Gal4* neurons in the pharyngeal region also blocked the PA-induced glucose feeding response (Fig. 1E).

We noticed that the two pharyngeal *Gr43a-Gal4* neurons are also labeled by a *npfr1-Gal4* driver, as evidenced by the overlapping GPF expression at the larval pharyngeal region (Fig. 2A-C). Consistent with this anatomical observation, functional knockdown of *npfr1* activity attenuated PA-aroused glucose feeding in *Gr43a-Gal4*/UAS-*npfr1*^*RNAi*^ fed larvae (Fig. 2D). Importantly, the fructose feeding activity of these fed larvae, which is independent of the odor stimulation, remained normal (Fig. 2E). In a control experiment, we showed that targeted lesioning of the two Gr43a-Gal4 neurons also blocked the fructose feeding response. Finally, using CaMPARI-based imaging^27^, we observed that the same pair of pharyngeal *npfr1-Gal4* neurons of fed larvae was activated after feeding in the fructose but not glucose medium) (Fig. S1). Together, our findings suggest that the Gr43a neurons mediate odor-aroused glucose feeding and odor-independent fructose feeding through two distinct intracellular signaling pathways.

**Figure 2.**
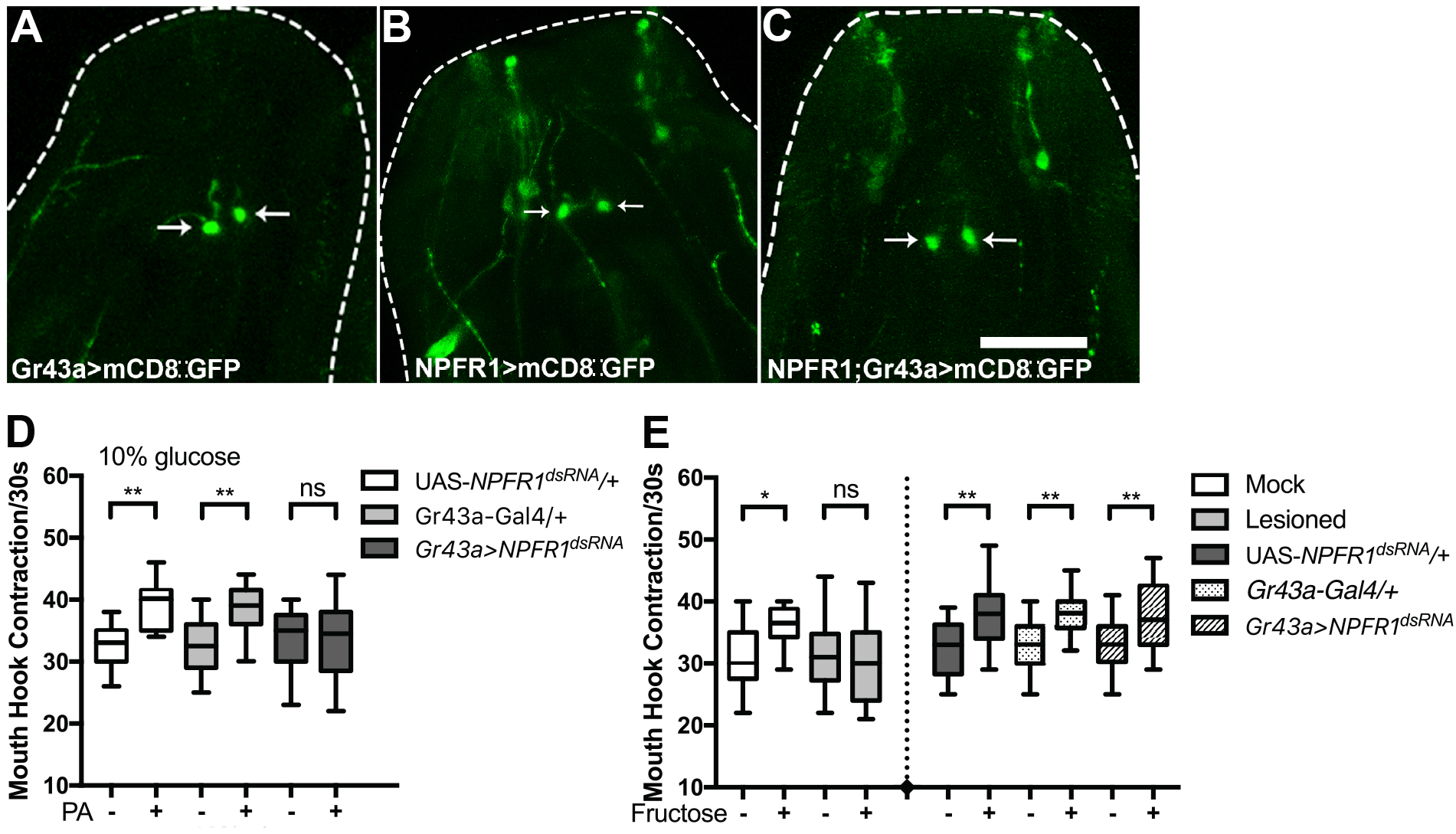
NPFR1 mediates integration of PA and glucose taste in the Gr43a neurons. **(A-C)** Expression of a GFP reporter driven by *Gr43a-Gal4* or *npfr1-Gal4* or both. A pair of *npfr1-Gal4* neurons overlaps with *Gr43a-Gal4* neurons in the pharyngeal region. n = 10-16. Scale bar = 100 μm. **(D)** Functional knockdown of npfr1 activity in *Gr43a-Gal4* neurons abolished PA-aroused glucose feeding. n = 20-27. Kruskal-Wallis test followed by Dunn’s multiple comparisons. ** P < 0.0001. **(E)** Fed larvae showed increased feeding in response to fructose, which was abolished by laser lesions in the two peripheral *Gr43a-Gal4* neurons. n = 12-20. In contrast, *npfr1* knockdown in the *Gr43a-Gal4* neurons failed to block the fructose feeding. n = 20-32. One-way ANOVA test followed by Sidak’s multiple comparisons. * P < 0.05, ** P < 0.001.

### NPF potentiation of Gr43a neurons for the adaptive response to glucose

Axons of the two pharyngeal Gr43a neurons are projected into the subesophageal zone (SEZ), a larval hindbrain-like region^23^ (also see Fig. 1B-D). To examine the roles of NPF and its receptor NPFR1 in the glucose response, a fluorescent voltage sensor, Ace2N-AA-mNeon, was used to analyze the membrane potential of the axon terminals of the Gr43a neurons in a live preparation of fed larvae^28^ (Fig. 3). As a positive control, 5% fructose stimulation of the larval preparation led to significantly increased depolarization at the axon terminals (Fig. 3A). In contrast, stimulation by 10% glucose or NPF alone failed to excite the axons. To our surprise, NPF and glucose co-stimulation led to a level of depolarization similar to that induced by fructose (Fig. 3B). These results reveal the presence of a previously uncharacterized smell-taste integration process at the axon terminals of two Gr43a gustatory neurons. Further, using CaMPARI-based imaging, we showed again that Gr43a neuronal response to glucose is NPF dependent (Fig. S2). Therefore, NPF signaling likely contributes to this process by potentiating Gr43a neuronal response to otherwise non-stimulating glucose.

**Figure 3.**
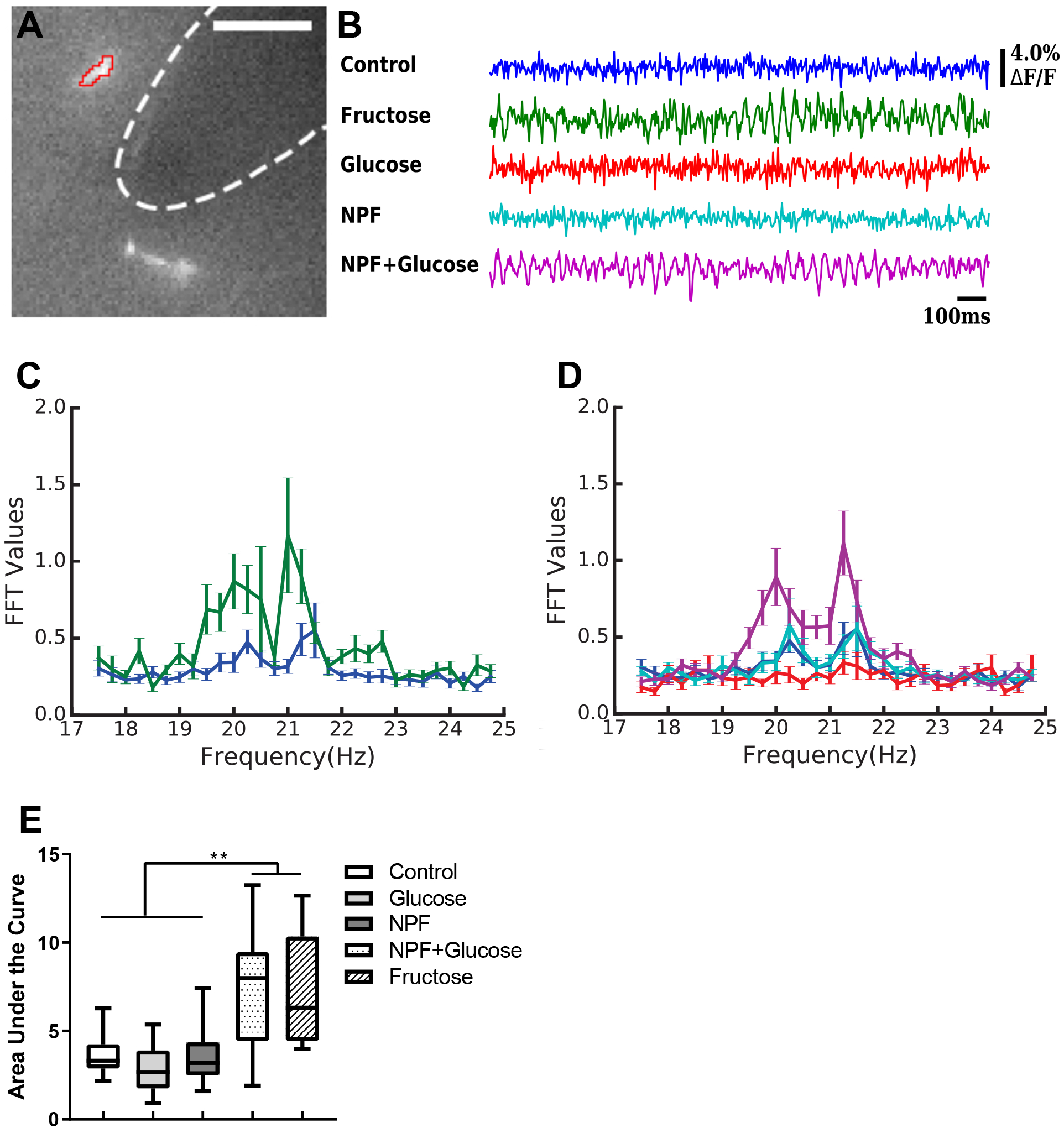
The Gr43a neurons displayed similar excitabilities in response to fructose or NPF/glucose co-stimulation. **(A)** An example of Gr43a axon terminal in the SEZ selected for mNeon-mediated imaging analyses. Scale bar = 25 um. Sample recordings of normalized Gr43a axon terminal membrane potential activities from: 5% fructose-stimulated intact larval CNS tissues and unstimulated controls; **(B)** recordings from glucose-, NPF- or glucose/NPF-stimulated tissues and unstimulated controls. NPF= 1 μM. (**C, D**) Quantification of the imaging data using fast Fourier transfer reveals significantly increased FFT values within the range of 19 to 23 Hz, when stimulated by both glucose and NPF or fructose alone(Pu et al., 2018). **(E)** Area under each curve between 19Hz to 22Hz was measured. n = 8-17. One-way ANOVA test followed by Dunnett’s multiple comparisons test. ** P < 0.01.

### Role of Gr43a neurons in adaptive feeding response to fatty acid

The glucose feeding of fed larvae is also enhanced by balsamic vinegar (BV) aroma^29^. Interestingly, unlike PA, this natural aroma also stimulates larval feeding of oleic acid media (Fig. 4A). To better understand the underlying integration processes, we first expressed Kir2.1 in the *Gr43a-Gal4* neurons. Again, silencing of the *Gr43a-Gal4* neurons completely abolished the BV-aroused feeding responses to oleic acid and glucose (Fig. 4B). In addition, functional knockdown of *npfr1* in the *Gr43a-Gal4* neurons also attenuated the BV-aroused feeding responses (Fig. 4C). We also performed a CaMPARI-based imaging analysis to investigate the role of Gr43a neurons in BV-aroused feeding behavior (Fig. 4D,E). In BV-aroused fed larvae, the Gr43a neurons were activated in response to oleic acid stimulation. However, the same neurons in unstimulated larvae failed to do so.

**Figure 4.**
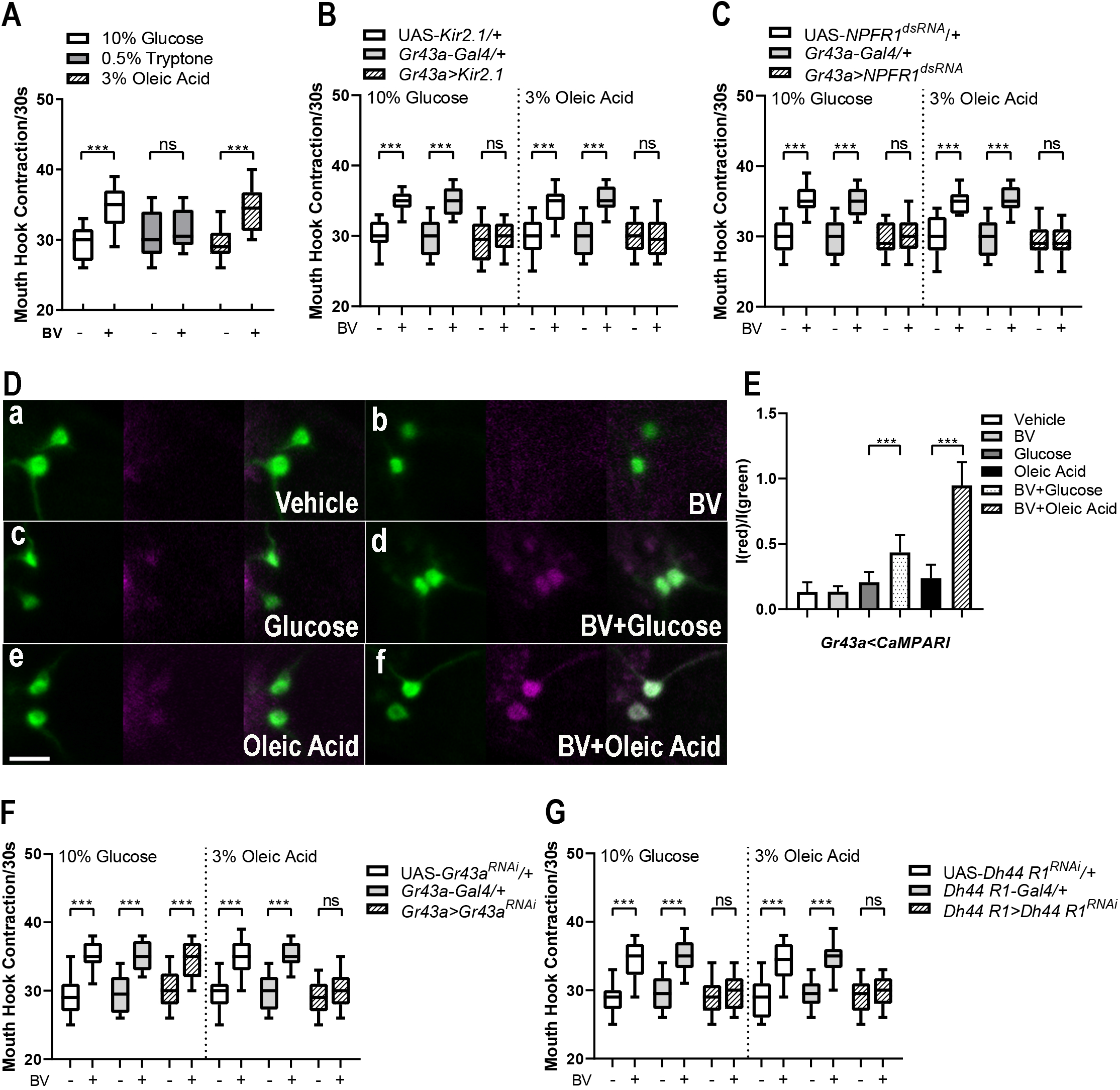
Analysis of balsamic vinegar aroma-aroused feeding and underlying gene and neural activities in fed larvae. **(A)** After stimulation by an effective dose of BV, fed larvae selectively increased feeding response to liquid media containing glucose or oleic acid. n = 12-16. **(B, C)** Kir2.1-mediated silencing of the Gr43a neurons or functional knockdown of *npfr1* activity abolished BV-aroused feeding of glucose or oleic acid. n =12-20. **(D)** CaMPARI-based imaging of the BV-aroused responses of two pharyngeal *Gr43a-Gal4* neurons in larvae fed with liquid media with or without 10% glucose or 3% oleic acid (b, d and f). In the absence of BV treatment, the Gr43a neurons in fed larvae showed no detectable responses to any of the three media (a, c and e). Scale bar = 40 mm. **(E)** Quantification of the CaMPARI signals in the *Gr43a-Gal4* neurons of fed larvae. n=11-17. **(F, G)** Functional knockdown of *Gr43a* activity in Gr43a-Gal4 neurons differentially affected larval feeding activities in media containing 10% glucose or 3% oleic acid, while *DH44R1* knockdown attenuated both feeding activities. n=15-20. One-way ANOVA followed by Dunnett’s multiple comparisons test. ***P<0.001

We also performed functional knockdown of additional genes using the *Gr43a-Gal4* driver. Gr43a encodes a fructose receptor, and is the only known sweet receptor in fly larvae^16,30^. Fed larvae expressing UAS-*Gr43a*^*RNAi*^ displayed normal glucose response stimulated by BV aroma, while their BV-aroused oleic acid feeding was abolished (Fig. 4F). Therefore, the Gr43a receptor appears to mediate larval sensing of both sugar and fatty acid. *DH44R1* encodes a receptor for a corticotropin releasing factor (CRF)-like neuropeptide (DH44), which is implicated in taste-independent sensing of nutritional sugars and their ingestion and digestion^31^. We also found that fed larvae expressing UAS-*DH44R1*^*RNAi*^, directed by *DH44R*I-*Gal4*, showed attenuated feeding responses to glucose and oleic acid (Fig. 4G). Therefore, the CRF-like receptor neurons may act downstream from the Gr43a neurons to define a feeding circuit for aroma-aroused appetite for palatable food.

### A G protein-mediated binary mechanism for tuning appetizing odor signals

A pair of NPF neurons (DM-NPF), each located at the dorsomedial region per brain hemisphere, defines a neural circuit module for appetizing odor perception and reward-motivated feeding. Each NPF neuron receives ascending food odor-evoked DA signals through the lateral horn (LH) per brain hemisphere, and relays descending NPF-encoded appetizing odor cues to the SEZ through their long projection axons^9^ (also see Fig. 6). The evidence described above suggests that the Gr43a neurons are potentiated by such NPF signals through activation of an otherwise silent NPFR1-dependent pathway for glucose sensing. Our previous study also showed that a D1-like DA receptor (Dop1R1) mechanism in the NPF neuron is responsible for selective conversion of a fraction of odor-evoked DA inputs to NPF outputs^9,18^. Therefore, we performed a CaMPARI-based imaging analysis to evaluate how Dop1R1 and related gene activities may contribute to PA-evoked NPF neuronal excitation. Fed larvae were first treated with appetizing or non-appetizing odor stimuli under the same conditions as those for behavioral assays, and their tissue preparations were harvested. In normal fed larvae, the NPF neurons displayed excitatory response to the stimulation by an appetizing dose of PA (7μl). However, stimulations of fed larvae using non-appetizing PA doses (e.g., 20 or 3.5μl) failed to do so (Fig. 5A). Furthermore, when Dop1R1-defieinct NPF neurons were tested, the effective dose of PA required to activate NPF signaling is upshifted from 7μl to 20μl PA. These results reveal that the inverted-U effects of odor stimuli can be observed at both behavioral and NPF neuronal levels^18^.

**Figure 5.**
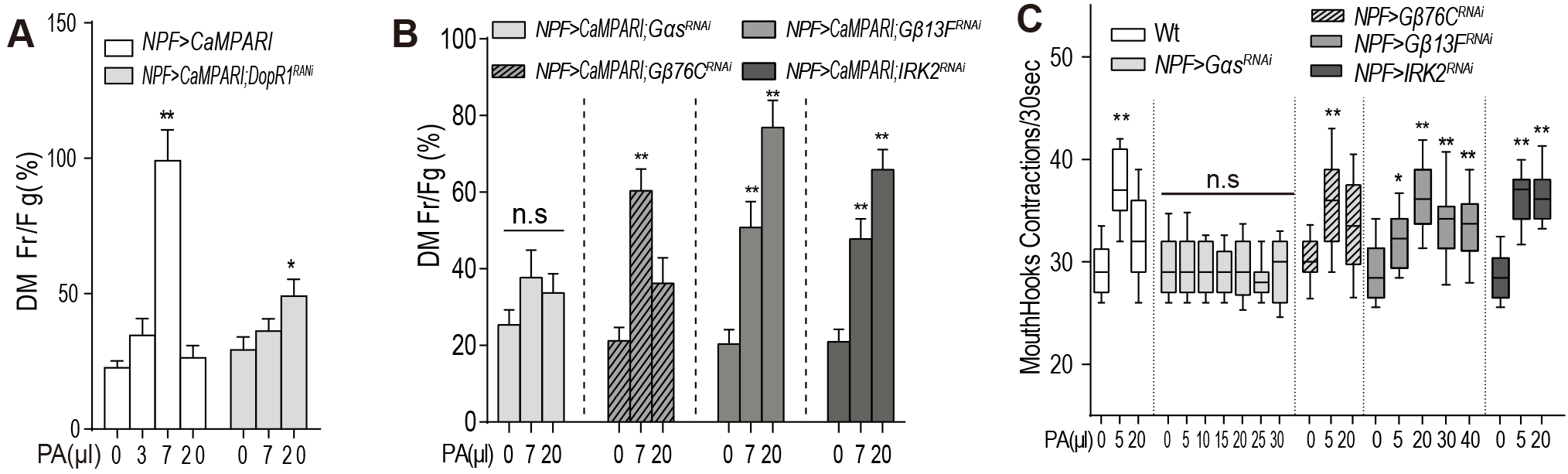
Genetic analysis of a G protein-gated binary tuning mechanism in NPF neurons. **(A)** CaMPARI-based imaging of the dorsomedial NPF neurons deficient for Dop1R1 showed a right-shift in the dose response by the NPF neurons, paralleling the previously observed right-shift in the dose-response curve of PA-aroused fed larvae^9^. n= 6-8. **(B)** CaMPARI-based imaging analyses of the NPF neurons: Gαs-deficient NPF neurons failed to display excitatory responses all doses tested, while *Gβ13F*- or *IRK2*-deficient NPF neurons were responsive to higher doses that are otherwise behaviorally ineffective as well as doses that are normally effective. n=8-10 **(C)** Behavioral analyses of PA-aroused feeding activities of fed larvae: Gαs-deficient larvae failed to respond to PA all doses tested; *Gb13F*- or *IRK2*-deficient larvae responded to PA at normally effective as well as at higher doses. n=14-18. One-way ANOVA followed by Dunnett’s multiple comparisons test. *P<0.05; **P<0.001.

**Figure 6.**
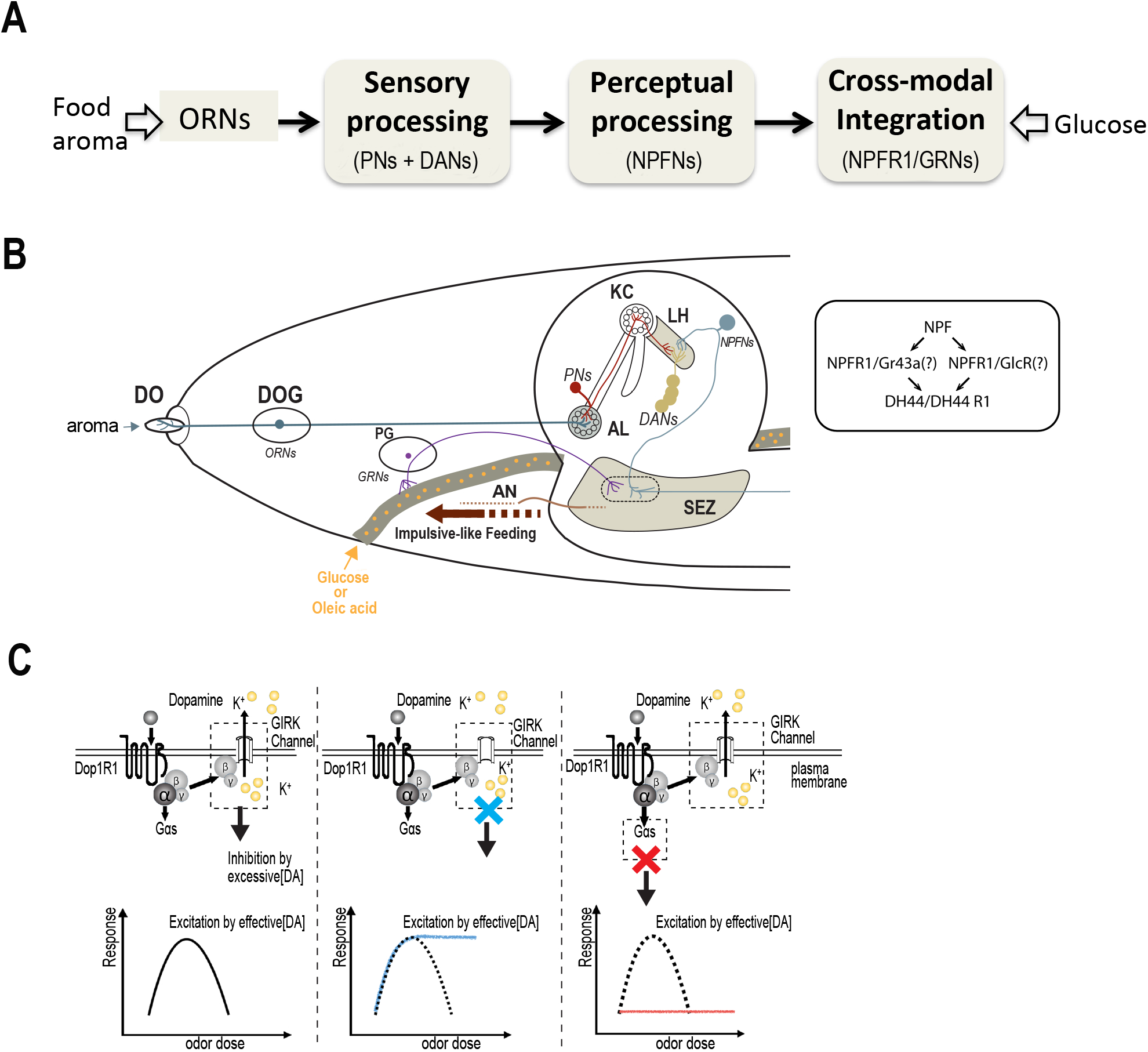
A schematic summary of the roles of DA, NPF and NPFR1 neurons in larval perception of congruent food aroma and glucose taste. **(A)** A schematic diagram showing a complete neuronal pathway for sensory and perceptual processing of congruent aroma and taste and integration of their neural representations. **(B)** Depiction of same neuronal pathway comprising DA, NPF and NPFR1-positive Gr43a neurons in the context of the larval nervous system. Only one of the two parallel pathways is shown. The molecular signaling pathway for glucose or oleic acid sensing by the Gr43a neurons is described in the box. **(C)** As shown above, he NPF neurons are the highest-order neurons in the pathway. They receive ascending presynaptic DA signals from the lateral horn, and relay descending NPF signals to the SEZ, where the axon terminals of NPFR1-positive Gr43a neurons are located. Left panel: a G protein-mediated binary mechanism tunes NPF neuronal response to DA in an inverted U manner; Middle: the deficiency in the Gβ13F/IRK2 pathway converts the inverted-U curve to a sigmoidal curve; Right: the deficiency in the Gαs pathway abolished NPF neuronal responses to DA at all odor doses tested. ORNs: olfactory receptor neurons; PNs: projection neurons; DANs: a subset of DA neurons named DL2-1 to 4^43^. NPFNs: NPF neurons at the dorsomedial region of the brain hemisphere; NPFR1/GRNs: NPFR1-positive Gr43a neurons; KC: Kenyon cells; AL: antenna lobes; LH: lateral horns; SEZ: the subesophageal zone; AN: antenna nerve; DO: dorsal organs; DOG: dorsal organ ganglia; PG: pharyngeal ganglia.

D1-type DA receptor is associated with the heterotrimeric G protein complex consisting of Gαs, Gβ, Gγ subunits^32^. Upon its activation by DA, the dissociated Gαs subunit and Gβγ complex each defines a separate effector pathway. We found that RNAi-mediated knockdown of Gαs activity blocked NPF neuronal response as well as behavioral response to PA at all doses tested (5 to 20 μl; Fig. 5B, C; Fig. S2). In parallel, we also functionally knocked down two Gβ genes. The NPF neurons expressing UAS-*Gβ13F*^*RNAi*^ displayed a normal response to 7μl PA as well as a strong excitatory response to a normally ineffective dose of PA (20μl). The Drosophila genome encodes three inwardly rectifying potassium channels (IRK1-3). We postulate that one or more of such IRKs may be a G protein-coupled IRK that could serve as a potential molecular target of the Gβγ complex^33–35^. To test this hypothesis, we functionally knocked down each of three channels (IRK1-3). Indeed, NPF neurons expressing UAS-*IRK2*^*RNAi*^, but not UAS-*IRK1*^*RNAi*^ or UAS-*IRK3*^*RNAi*^, showed altered cellular responses to odor stimulation, similar to UAS-*Gβ13F*^*RNAi*^-expressing NPF neurons (Fig. 5B).

To corroborate the cellular imaging results, we also analyzed the PA-evoked glucose feeding of *npf-*GAL4/UAS-*Gβ13F*^*RNAi*^ and *npf-*GAL4/UAS-*IRK2*^*RNAi*^ fed larvae. As expected, both fed larvae showed broader response profiles at all PA doses tested (from 5 to 40μl). In addition, functional knockdown of two other IRK genes, IRK1 and IRK3, had no effect on the feeding behavior of fed larvae (Fig. 5C, Fig. S3). Therefore, both imaging and behavioral analyses suggest that a binary genetic mechanism involving a Gαs and a Gβ13F/IRK2 pathway mediates the precision tuning of Dop1R1-gated NPF neuronal responses to odor stimuli of various doses. When the Gβ13F/IRK2 pathway is deficient, the effective range of DA-encoded odor inputs is greatly expanded, converting the inverted U dose-response curve to the sigmoidal shape in these larvae.

## Discussion

We have provided examples how pairing of food aroma (e.g., a banana-like or balsamic vinegar scent) with glucose or oleic acid taste drastically enhanced appetite in well-nourished *Drosophila* larvae. Using these behavioral paradigms, we investigated the molecular and cellular underpinnings of the appetizing effects of the congruent aroma-taste pairs. One of the key findings is that the congruency of an aroma and glucose or oleic acid taste requires a neural circuitry involving two peripheral Gr43a gustatory neurons projecting to the hindbrain-like SEZ as well as two brain NPF neurons that modulate the Gr43a neurons (Fig. 6A). In addition, the appetizing effect of the aroma and food taste requires a cross-modal integration process that takes place at the axon terminals of the Gr43a neurons, and is mediated by a NPF/NPFR1-mediated signaling mechanism that potentiates Gr43a neuronal response to otherwise non-stimulating glucose and oleic acid (Fig. 6B). We have also identified a G protein-mediated tuning mechanism in the NPF neurons that selectively converts appetizing odor-evoked DA signals to descending NPF-encoded appetitive signals relayed to the SEZ (Fig. 6C). Our finding of congruent aroma-induced expansion of taste neuron response profiles suggests that food taste sensation is significantly more complex that commonly recognized.

### NPF-induced expansion of Gr43a neuronal response profiles

It remains unclear how NPF activation of its receptors in the axonal terminals of the Gr43a neurons enables the Gr43a neuronal response to otherwise non-stimulating food taste. NPF signaling may act in two different ways. The first possibility is that NPF signaling may drastically increase the reactivity of a gustatory receptor pathway to its ligand. For example, it could rapidly increase the number of available receptors thorough transfer of internally stored receptors to the dendritic surface. We have shown that the gustatory receptor Gr43a is required for Gr43a neuronal response to oleic acid as well as fructose. However, unlike the fructose response, the response to oleic acid also depends on potentiation of the Gr43a neurons by congruent aroma-evoked NPFR1 activation. Therefor, it would be interesting to determine whether BV stimulation may significantly increase the number of Gr43a receptors on dendritic surface.

The second possibility is that NPF signaling may lead to the activation of dormant gustatory receptors in the Gr43a neurons (e.g., thorough receptor complex formation or chemical modification). We have shown that aroma-stimulated fed larvae displayed virtually identical feeding responses to liquid media containing a broad concentration range of glucose from 2.5 to10%^9^. These results are consistent with the notion that NPFR1 activation may turn on a yet uncharacterized glucose receptor that is functionally silent in the Gr43a neurons. At present, while multiple gustatory receptor (Gr) genes for bitter compounds are expressed in fly larvae, Gr43a remains the only known sweet receptor^16,24^. Glucose sensing by the Gr43a neurons may involve other classes of receptors such ionotropic receptors (Ir)^35–37^. It also remains to be determined whether NPFR1, a G-protein coupled receptor, directly or indirectly mediates glucose binding.

*DH44R1* encodes a receptor for a corticotropin releasing factor (CRF)-like neuropeptide (DH44), which is implicated in taste-independent sensing of nutritional sugars and their ingestion and digestion^30^. However, our study suggests that the CRF-like signaling mechanism, mediated by DH44R1, plays more diverse roles in feeding responses, including taste-dependent intake of different energy-dense foods. We propose that DH44R1 neurons define a regulatory module that acts downstream from the Gr43a neuron-mediated sensory/integration module (Fig. 6B). Together, they contribute to a feeding circuit for aroma-aroused feeding of palatable food.

### A second contribution of NPF neurons to the aroma-glucose congruency

The paired NPF neurons, located at the dorsomedial region of the brain, are central to the larval food reward system^29^. Each of them receives ascending DA-encoded olfactory inputs from the lateral horn ipsilaterally. Genetic silencing of the two NPF neurons blocked the odor-aroused glucose feeding in fed larvae, while their genetic activation triggered glucose overeating in the absence of odor stimulation. In this study, we show that these NPF neurons also make a second contribution to the aroma-glucose congruency by releasing NPF signals in an appetizing odor-dependent manner. CaMPARI-based imaging revealed that the NPF neurons are activated by appetizing odor stimuli in behaving fed larvae, but they failed to respond to behaviorally ineffective PA stimuli that are either too strong or too weak. The imaging assay also showed that *Dop1R1-*deficient NPF neurons displayed a right-shift in their dose response curve (Fig. 5A). This finding suggests that a DA/Dop1R1 mechanism regulates the response pattern of the NPF neurons to appetizing odor stimuli, which in turn shapes the inverted U dose-response curve of odor-stimulated fed larvae^29^.

### A G protein-mediated binary mechanism in tuning NPF neuronal signaling

So far, our findings show that Dop1R1-gated NPF neuronal signaling represents a crucial regulatory step for pairing glucose taste with congruent PA stimuli. Using both functional imaging and behavioral assays, we have identified two separate molecular signaling pathways that together define a two-layered strategy for precision tuning of the NPF neuronal responses to appetizing odor-evoked DA inputs. First, attenuation of the Gβ13F or IRK2 activity greatly broadens the width of the effective dose range of DA signals, as evidenced by the change of the dose-response curve from an inverted U shape to a sigmoidal shape (Fig. 6C). These results suggest that Gβ13F/IRK2-mediated pathway is dispensable when DA inputs are not excessively high, and it sets an upper limit of the optimum effective range of odor-induced DA signals by silencing NPF neuronal responses to odor stimuli whose strengths are above the upper limit. The second Gαs-mediated pathway provides a default mechanism essential for the excitatory responses of NPF neurons to any odor stimuli above minimum threshold strength. However, the effect of this Gαs pathway is dominantly inhibited once the Gβ13F/IRK2 pathway is activated by an excessively strong odor stimulus.

### Conserved neural substrates and mechanisms in aroma-taste congruency

Food flavor pairing is widely used to enhance the appetitive value of meals in humans and animals alike. We have provided evidence that in fed larvae, at least two different neural processes underlying food aroma-taste congruency involve conserved neural substrates and mechanisms. For example, the conserved DA and NPY-like systems play crucial roles in appetizing odor perception by selectively converting a fraction of food odor stimuli to NPF-encoded rewarding cues that are relayed to the taste-processing center. Similarly, the inverted-U effects of DA on cognitive functions have been widely observed in humans and other animals^38,39^. In the prefrontal cortex of mammals, an optimum level of D1-type DA receptor activity is crucial for spatial working memory, while its signaling at levels that are too low or too high leads to impaired working memory^40–42^. In addition, the neural process for selective pairing of food aroma and glucose taste also involves NPY-like and CRF-like systems. Based on these findings, we propose that a variety of conserved neuropeptide systems could be activated by appetizing olfactory or other sensory stimuli, thereby leading to alteration of the response profiles of different subsets of gustatory neurons for food flavor recognition. In the mammalian brain, the stimuli of food smell and taste appear to interact within the gustatory cortex^5–8^. Therefore, it will be interesting to determine whether homologous molecular and neural activities are involved in higher-order neurons that modulate gustatory neurons for food tasting.

## Methods

### Fly Stocks and Larval Growth

All flies are in the w1118 background. Larvae were reared at 25°C, and early third instars (~74 hr after egg laying, AEL) were fed before behavioral experiments as previously described^9^. The transgenic flies include *npfr1-Gal4* (Wang et al., 2013), *Gr43a-GAL4* (BL57636), *UAS-mCD8∷GFP^9^, UAS-Kir2.1*^44^, UAS-*Gr43a*^*RNAi*^, UAS-Gαs^RNAi^ (BL29576), UAS-Gβ76^CRNAi^ (BL28507), UAS-Gβ13F^RNAi^ (BL35041), UAS-Ace2N-AA-*mNeon* (BL64317), UAS-*CaMPARI* (BL58761), UAS-*IRK2*^*RNAi*^ (BL41981), UAS-*IRK1*^*RNAi*^ (BL42644) and UAS*-IRK3*^*RNAi*^ (BL26720) were obtained from Bloomington stock center. UAS-*Dop1R1*^*RNAi*^ (V107058) was obtained from the Vienna Drosophila RNAi Center.

### Behavioral assays

Quantification of mouth hook contraction rate in liquid food was performed as previously described^44^. Odor stimulation of fly larvae was performed using a published protocol^9^. After rinsing with water, larvae were tested for their feeding responses. Feeding media include agar paste (US Biological, A0940) containing 10% glucose, 0.5% Tryptone (Becton-Dickinson, 628200) or 3% oleic acid (Sigma-Aldrich, 112-80-1). All behavioral assays were performed under blind conditions.

### Laser lesion

Laser lesion was performed using a previously published protocol with slight modification^22^. Early second-instar larvae (52 h after egg laying) were transferred to 150μl double-distilled H2O on a microscope slide, and placed into the anesthetization chamber (90-mm Petri) with ether for 2.5 min. The laser beam was focused on individual Gr43a nuclei as a burst of 20 shots at a rate of 3 Hz. The mock group contains larvae that went through all the treatments except laser lesion. After the laser treatment, the larvae recovered on fresh apple juice plates with yeast paste for 24 h before the assay.

### CaMPARI imaging

The odor treatment conditions for CaMPARI imaging are identical to those for odor-aroused feeding behavioral assays. After odor stimulation, larvae were irradiated with PC light 405 nm LED array (200 mW/cm^2^, Loctite) for 5s. The treated larval CNS tissues were processed as previously described(Pu et al., 2018). The treated larval preparation was imaged using a Zeiss LSM 510 confocal microscope.

### mNeon imaging and data processing

For mNeon Imaging, a published protocol was followed with minor modifications^29^. Briefly, immediately after dissection, the larval CNS preparation was incubated in *Drosophila* PBS, and imaged under 40X water immersion lens using Zeiss Axio Examiner. NeuroCCD-SM camera. Effective and ineffective odor vapors were prepared by fumigating a sealed 24L foam box with 150 or 800μl PA for 2 hours, respectively. Odor was continuously delivered to larval head region by pumping at the rate of 0.36L/min. Turbo-SM software (RedShirt Imaging) was used for recording and data processing. Images were captured at a frame rate of 250 Hz, and exposure time is 10ms. 1000 frames were collected over a 4s period. Standard deviation and fast Fourier transform were obtained using normalized data.

### Statistical analysis

Statistical analyses were performed using Kruskal-Wallis test followed by Dunn’s multiple comparisons test or one-way ANOVA followed by Sidak’s or Dunnett’s multiple comparisons test. Data normality was tested by Shapiro-Wilk normality test.

## Acknowledgements

We thank Melissa M.M. Palombo for technical assistance. This work is supported by grants from the U.S. National Institutes of Health (DK102209 and DC013333) to P.S.

## Author Contributions

Y.W. Z. and Y.P. have performed imaging and behavioral assays and helped with writing the manuscript. Y. Z. and H. C. performed anatomical and functional analyses of DH44. P.S. supervised the work and wrote the manuscript.

---

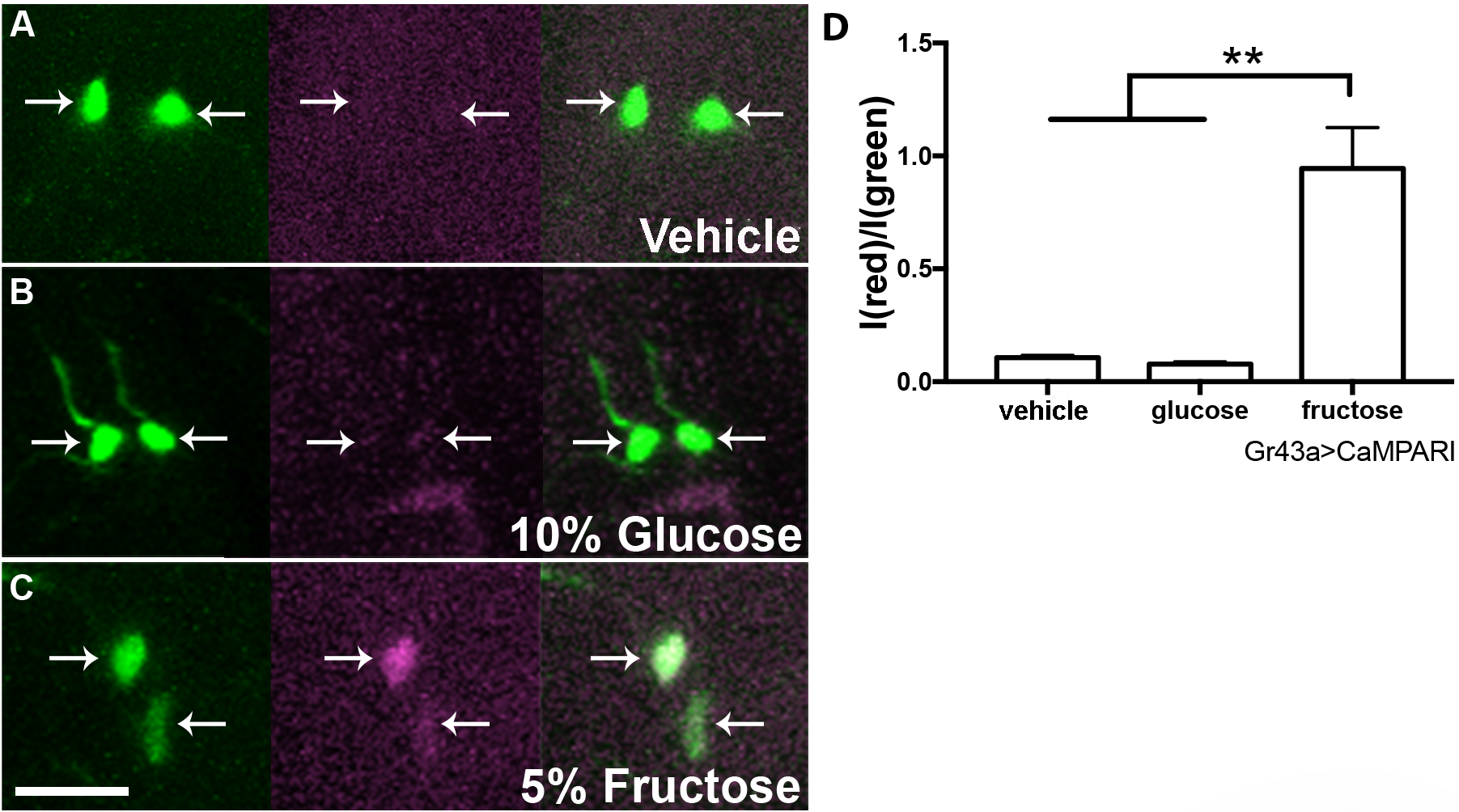

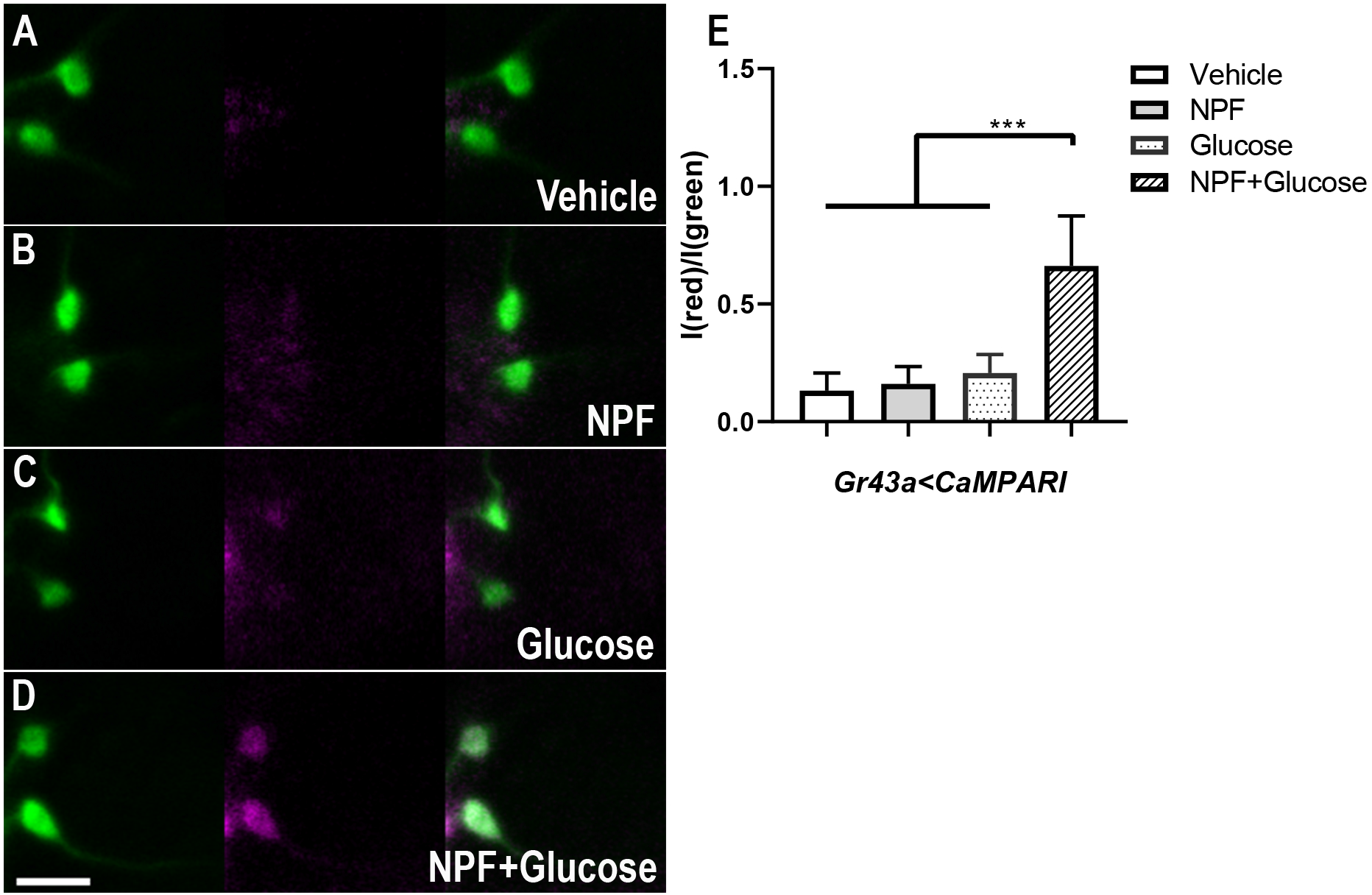

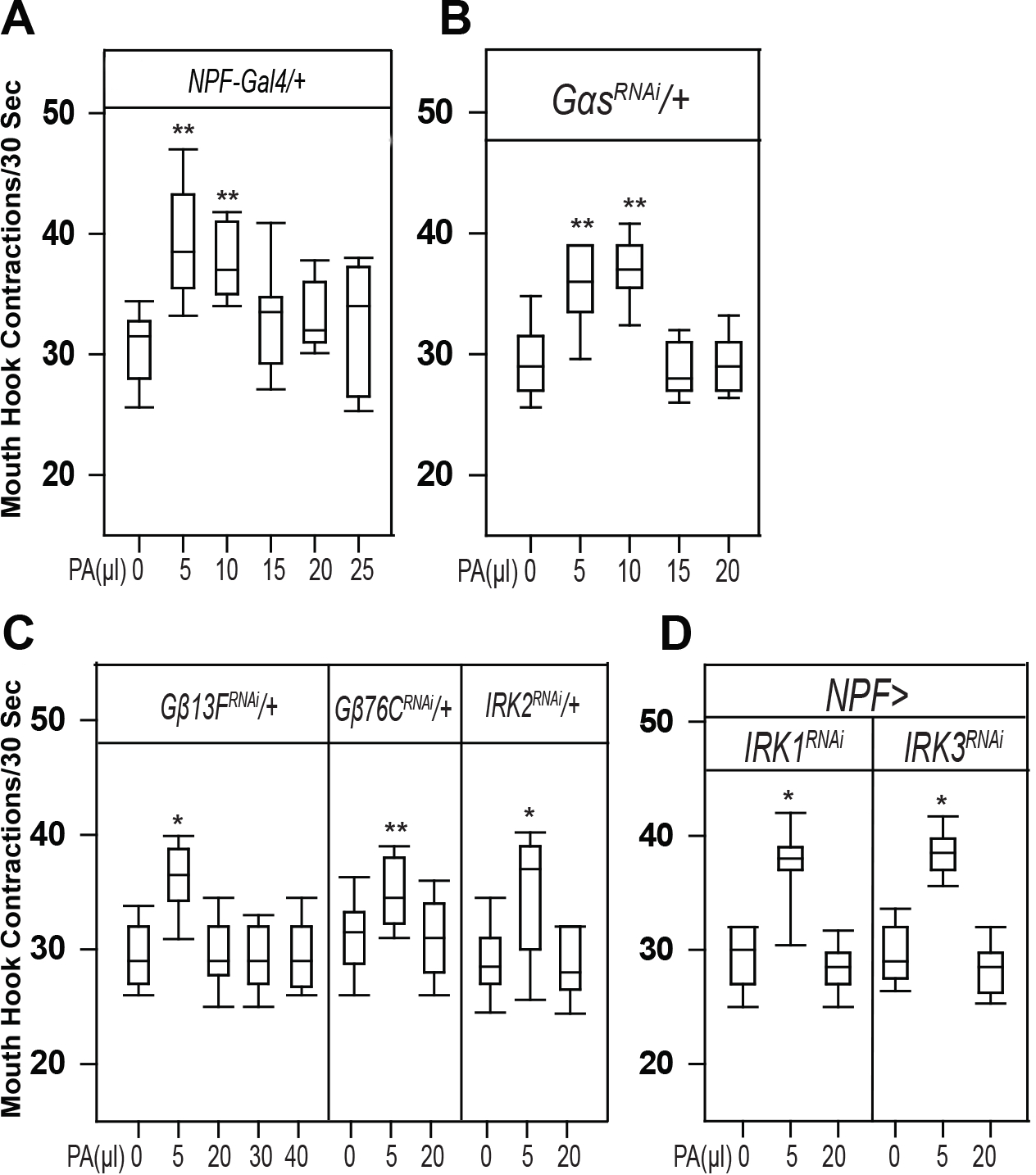

